# Macrophages induce inflammation by efferocytosis of apoptotic prostate cancer cells via HIF-1α stabilization

**DOI:** 10.1101/2021.09.02.458687

**Authors:** Veronica Mendoza-Reinoso, Patricia M. Schnepp, Dah Youn Baek, John R. Rubin, Ernestina Schipani, Evan T. Keller, Laurie K. McCauley, Hernan Roca

## Abstract

Clearance of apoptotic cancer cells by macrophages, known as efferocytosis, fuels the bone-metastatic growth of prostate cancer cells via pro-inflammatory and immunosuppressive processes. However, the exact molecular mechanisms remain unclear. In this study, single-cell transcriptomics of bone marrow macrophages undergoing efferocytosis of apoptotic prostate cancer cells revealed a significant enrichment of a cellular response to hypoxia. Here we show that efferocytic macrophages promote HIF-1α stabilization under normoxic conditions through interaction with phosphorylated STAT3. Inflammatory cytokine gene expression analysis of efferocytic HIF-1α-mutant macrophages revealed a reduced expression of the pro-tumorigenic *Mif*. Furthermore, stabilization of HIF-1α using the HIF-prolyl-hydroxylase inhibitor, Roxadustat, enhanced MIF expression in macrophages. Finally, macrophages treated with recombinant MIF protein activated NF-κB (p65) signaling and increased the expression of pro-inflammatory cytokines. Altogether, these findings suggest that the clearance of apoptotic cancer cells by tumor-associated macrophages triggers p-STAT3/HIF-1α/MIF signaling to enhance tumor-promoting inflammation in bone, suggesting this axis as a target for metastatic prostate cancer.

## Introduction

The process of clearing of apoptotic cancer cells by macrophages, known as efferocytosis, commonly occurs during tumor progression and fuels the bone-metastatic growth of cancer cells, via subsequent pro-inflammatory and immunosuppressive activity (Graham *et al*, 2014; Lecoultre *et al*, 2020; Roca & McCauley, 2018; Stanford *et al*, 2014). Our previous published work reported that bone marrow macrophage-dependent efferocytosis of apoptotic prostate cancer cells supported skeletal tumor growth through the secretion of pro-inflammatory cytokines resulting in an immunosuppressive response (Mendoza-Reinoso *et al*, 2020; Roca *et al*, 2018). Recently, a single-cell RNA sequencing study reported that peritoneal macrophage efferocytosis of apoptotic T cells displayed a heterogeneous transcriptional activity including genes associated with predisposition for efferocytosis, macrophage differentiation, locomotion and inflammation (Lantz *et al*, 2020). However, the precise molecular mechanisms involved in bone marrow macrophage response to efferocytosis of apoptotic cancer cells remains to be elucidated.

The majority of solid tumors present areas of permanent or transient hypoxia due to poor vascularization and blood supply (Pouyssegur *et al*, 2006). Hypoxic conditions activate hypoxia inducible factor (HIF) signaling which has a crucial role in pro-tumorigenic inflammatory processes via cytokine secretion, reactive oxygen species (ROS) production and angiogenesis (Triner & Shah, 2016). HIFs are heterodimers consisting of an oxygen labile alpha (α) subunit and a stable beta (β) subunit. There are three isoforms of HIF-α, including HIF-1α, HIF-2α (EPAS1) and HIF-3α (IPAS) (Kaelin & Ratcliffe, 2008). HIF-1α upregulates glycolytic genes such as phosphoglycerate kinase (PGK) and lactate dehydrogenase A (LDHA) whereas HIF-2α induces the expression of genes related to oxygen supply improvement in hypoxic regions such as erythropoietin (EPO) (Hu *et al*, 2003). HIF-1α has been identified as a key regulator of proliferative, invasive and immunosuppressive mechanisms that favor tumor progression (Engel *et al*, 2017; Hatfield *et al*, 2019). Under hypoxic conditions, HIF-1α hydroxylation by prolyl hydroxylase is reduced. This inhibits the HIF-1α/Von Hippel-Lindau (VHL) interaction and consequent HIF-1α degradation by ubiquitin E3 ligase complex (Jaakkola *et al*, 2001). Therefore, HIF-1α is stabilized in the cytosol and translocated to the nucleus to promote the transcription of multiple target genes (Semenza, 2011). HIF-1α is strikingly upregulated under hypoxic conditions, however, HIF-1α can also be regulated at transcriptional, translational and post-translational levels under normoxic conditions (Hayashi *et al*, 2019). HIF-1α is expressed and stabilized in immune cells via hypoxia or other factors such as inflammation, cancer and infectious microorganisms (Blouin *et al*, 2004; Hartmann *et al*, 2008; Peyssonnaux *et al*, 2005). HIF-1α is crucial for myeloid cell-mediated inflammation (Cramer *et al*, 2003) and it has been demonstrated that tumor-associated macrophages (TAMs) also express HIF-2α under hypoxic conditions (Imtiyaz *et al*, 2010; Talks *et al*, 2000). Various studies have shown a relationship between HIF-1α induction and STAT3 activation at post-translational and transcriptional levels (Gray *et al*, 2005; Jung *et al*, 2008; Niu *et al*, 2008; Xu *et al*, 2005).

Previous reports have shown that low oxygen concentrations in tumors promote the secretion of cytokines and chemokines that recruit pro-tumorigenic Tregs, tumor-associated macrophages, neutrophils, B cells, and myeloid-derived suppressor cells (MDSCs) to support tumor growth (Blaisdell *et al*, 2015; Du *et al*, 2008; Facciabene *et al*, 2011; Zhu *et al*, 2014). One of these cytokines is macrophage migration factor (MIF), which is a direct target gene of HIF-1α (Winner *et al*, 2007) and a hypoxia-induced gene in colon and breast cancer cells (Larsen *et al*, 2008; Yao *et al*, 2005). MIF acts as an autocrine or paracrine cytokine, it is upregulated in several types of cancer (Mawhinney *et al*, 2015; Tomiyasu *et al*, 2002; Wilson *et al*, 2005) and its expression correlates with disease malignancy and invasiveness (Lippitz, 2013). Studies consistently demonstrated that MIF signals primarily through CD74 in association with CD44, CXCR2, CXCR4, and CXCR7 to activate the ERK MAP kinase cascade (Leng *et al*, 2003; Shi *et al*, 2006; Tarnowski *et al*, 2010). Finally, MIF signaling induces the activation and secretion of pro-tumorigenic cytokines to support tumor growth (Bach *et al*, 2008; Bucala & Donnelly, 2007).

Using single-cell transcriptomic sequencing, we investigated the signature changes in bone marrow macrophage gene expression during efferocytosis of apoptotic prostate cancer cells. We found that macrophages engulfing apoptotic prostate cancer cells promoted HIF-1α stability by its interaction with phosphorylated STAT3 and induced the expression of the pro-inflammatory cytokine MIF. Thus, p-STAT3/HIF1α/MIF signaling in tumor-associated macrophages may have a pro-tumorigenic effect in the bone marrow tumor microenvironment that contributes to skeletal metastasis and can be used to target preventive and therapeutic approaches.

## Results

### Single cell analyses of macrophages engulfing apoptotic prostate cancer cells show a distinct transcriptional signature and the activation of hypoxia-related genes

Published findings suggest that macrophages induce distinctive tumor-promoting signaling in response to efferocytosis of apoptotic cancer cells (Mendoza-Reinoso *et al*., 2020; Roca *et al*., 2018; Soki *et al*, 2014; Stanford *et al*., 2014). However, the mechanisms that govern these specific responses in connection to tumor acceleration are not completely understood. To further investigate the efferocytosis-mediated signaling in bone macrophages, primary bone marrow macrophages from immunocompetent C57BL/6J mice were co-cultured with CFSE^+^ (pre-labeled) apoptotic prostate cancer RM1 cells. Macrophages engulfing these apoptotic cancer cells (efferocytic) were compared to non-engulfing (non-efferocytic) macrophages by single-cell RNA sequencing upon sorting by flow cytometry (Figure 1A). After identification of high quality sequenced single cells (Table 1, Supplementary data), UMAP (Becht *et al*, 2018) was applied for dimension reduction of single cell analysis and visualization of the transcriptional data for efferocytic and non-efferocytic macrophages. As shown in Figure 1B the cell distribution in two UMAP projections depicts different cluster enrichment in efferocytic vs. non-efferocytic macrophages from two independent experiments (Exp. 1 and Exp. 2). For example, efferocytic cells demonstrate enriched in cluster in the direction of increased UMAP-2-projection while the opposite is observed in non-efferocytic macrophages as shown in the split visualization of these cells (Figure 1B). These results correlate with distinct transcriptional heatmaps in efferocytic relative to non-efferocytic macrophages (Figure 1C) where the great majority of differentially expressed genes (DEG’s) significantly changed in the same direction (3277 vs 482) in both experiments as analyzed in the Venn diagram in Figure 1D. DEG’s commonly upregulated in efferocytic macrophages in both experiments were further processed using the PANTHER analysis (Mi *et al*, 2013; Thomas *et al*, 2003) and gene ontology database (Ashburner *et al*, 2000; Gene Ontology, 2021) to identify the relevant biological pathways. Among the significantly enriched GO-biological processes identified we found pathways related to the innate immune system and wound healing responses which are related to phagocytosis of apoptotic cells, inflammation, and regeneration (Figure 1E, list of enriched GO terms in Table 1, Supplementary data) (Koh & DiPietro, 2011; Minutti *et al*, 2017). Intriguingly, biological processes related to hypoxia were identified even though these experiments were performed under normoxia (normal oxygen conditions), which suggests that efferocytosis mediated activation and upregulation of factors directly related to cellular hypoxia is independent of the oxygen concentration (Figure 1E and Figure 1F).

**Figure 1.**
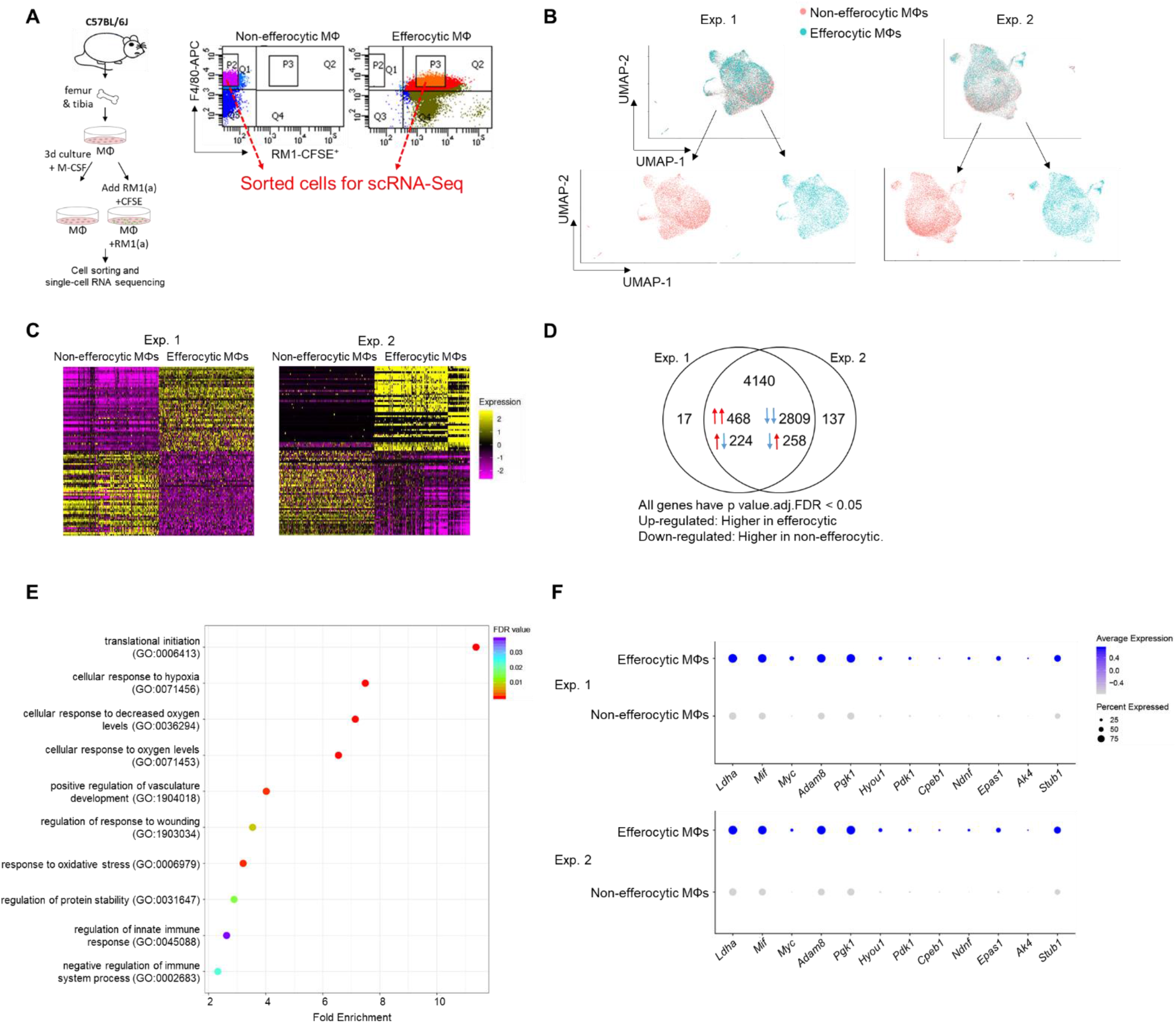
Single-cell experiments comparing efferocytic macrophages (engulfing apoptotic cancer cells) vs control (non-engulfing macrophages) **A.** Bone marrow-derived macrophages were isolated from 4 week old C57BL/6J mice and co-cultured with apoptotic RM1(a) cells for 16-18 hours. Efferocytic (F4/80^+^CFSE^+^) and control macrophages (F4/80^+^) were sorted and used for single-cell libraries followed by single-cell RNA sequencing. **B.** UMAP of all cells, blue represents efferocytic macrophages and red represents non-efferocytic macrophages. **C.** Heatmap of top differential expressed genes in either experiment. **D.** Venn Diagram of overlap of all differentially expressed genes between both experiments. **E.** Cleveland plot of top enrichment pathways in efferocytic macrophages. **F.** Dot plots of hypoxia-related genes in both experiments. Size relates to the percentage of macrophage cells that expressed each gene. Color denotes the average expression for each gene across all expressing cells. Additional results are shown in Supplementary data – Table 1, 2 and 3.

**Table 1.** Features of the data included in the single-cell RNA sequencing analysis.

| Sample | # of cells | # of unique genes per cell | # of transcripts per cell | Ave. mitochondrial % |
| --- | --- | --- | --- | --- |
| Non-efferocytic MΦ (1) | 7,813 | 3,029 | 12,837 | 3.39 |
| Efferocytic MΦ (1) | 6,262 | 3,726 | 19,768 | 3.18 |
| Non-efferocytic MΦ (2) | 10,612 | 2,909 | 11,326 | 2.89 |
| Efferocytic MΦ (2) | 7,984 | 3,800 | 17,984 | 3.27 |

### Efferocytosis of apoptotic cancer cells stabilizes HIF-1α in bone macrophages which is mediated by the activation of STAT3

Single cell analysis of efferocytic macrophages identified upregulated molecules associated with gene ontology (GO) term of cellular hypoxia (Figure 1). To investigate these findings in the overall macrophage population, co-cultures of bone macrophages with apoptotic prostate cancer RM1 cells (workflow Figure 1A) were analyzed. Selected hypoxia-GO associated genes showing significant upregulation by sc-RNAseq were investigated by RT-qPCR from total RNA isolated from co-cultures of independent bone macrophage populations. Fold changes in efferocytic relative to control macrophages were calculated and plotted in Figure 2A. The majority of analyzed genes showed a significant mRNA increase in efferocytic macrophages relative to control in correlation with the single cell results. Some of these molecules are crucial components of glycolysis including *Pdk1*, *Pgk1*, and *Ldha* and are known targets of hypoxia inducible factor 1a (HIF-1α), a master transcriptional regulator of cellular response to hypoxia that promotes metabolic switch to glycolysis (Niu *et al*., 2008; Palazon *et al*, 2014). *Hif1a* was upregulated in the overall population of efferocytic macrophages (although it was not identified as an upregulated gene in the single-cell experiments). Contrary to the results observed by single-cell experiments, the hypoxia-inducible transcription factor *Epas1* (HIF-2α) (Imtiyaz *et al*., 2010) showed a small but significant decrease in mRNA expression by efferocytosis.

**Figure 2.**
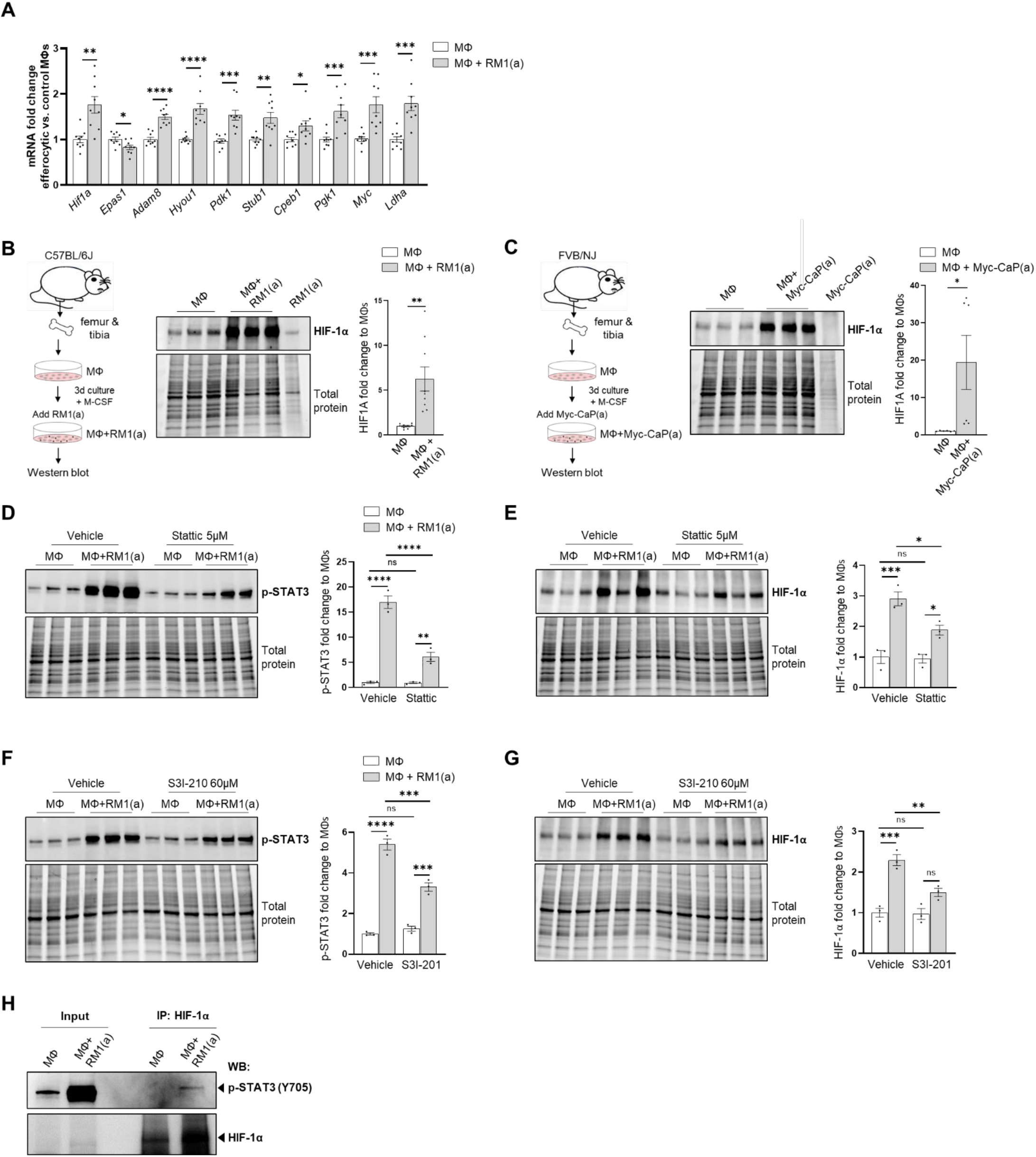
Macrophage efferocytosis of apoptotic cancer cells promotes HIF-1α stabilization through STAT3 activation. Bone marrow-derived macrophages were isolated from 4 week old C57BL/6J or FVB/NJ mice and co-cultured with apoptotic RM1(a) or Myc-CaP(a) cells for 16-18 hours. **A.** mRNAs isolated from efferocytic and control macrophages were analyzed by quantitative PCR (qPCR) for a set of genes involved in the cellular response to hypoxia (n=9). **B. & C.** Protein lysates from C57BL/6J (n=9) and FVB/NJ (n=6) efferocytic macrophages were analyzed by western blot using HIF-1α antibody. Protein lysates from C57BL/6J efferocytic and control macrophages (n=3) treated with 5μM Stattic and 60μM S3I-201 STAT3 inhibitors for 2 hours were analyzed by western blot using **D. & F.** Phospho-STAT3 antibody and **E. & G.** HIF-1α antibody. **H.** Cell lysates from efferocytic and non-efferocytic macrophages were immunoprecipitated with anti-HIF-1α antibody and immunoblotted with phospho-STAT3 antibody. Data plotted are mean ± SEM, *p < 0.05, **p < 0.01, ***p < 0.001, ****p < 0.0001, ns=not significant (Ordinary one-way ANOVA; Tukey’s multiple-comparisons test and unpaired t-test). Additional results are shown in Figure 2–Figure supplement 1 and 2.

Because HIF-1α is largely regulated post-transcriptionally resulting in a protein targeted for degradation under normoxic conditions (Ivan *et al*, 2001; Jaakkola *et al*., 2001), HIF-1α protein was further evaluated by Western blot. As shown in Figure 2B, Western blot analysis of efferocytic macrophages (co-cultured for 16-18 hours with apoptotic RM1 cells) evidenced a significant increase of HIF-1α induced by efferocytosis. These findings were further corroborated in efferocytic bone marrow derived macrophages co-cultured with murine prostate cancer Myc-CaP cells, which share several molecular characteristics of human prostate cancer (Dudzinski *et al*, 2019; Ellwood-Yen *et al*, 2003). As Myc-CaP cancer cells were obtained from FVB/NJ mice, the primary macrophages were obtained from the bone marrow of the same strain. Similar results were observed using this model (Figure 2C). These results suggest that HIF-1α is stabilized in macrophages engulfing apoptotic cancer cells.

The potential mechanism inducing HIF-1α stabilization was further investigated. Previous findings suggested that one potential mechanism leading to HIF-1α stabilization is via interaction with activated (phosphorylated) STAT3 (Gray *et al*., 2005; Jung *et al*., 2008; Li *et al*, 2019; Xu *et al*., 2005). Since STAT3 activation is sustained in efferocytic macrophages and considered a hallmark macrophage response to engulfing apoptotic cancer cells it was hypothesized that STAT3 activation is critical in HIF-1α stabilization by efferocytosis. We investigated this using two well characterized STAT3 inhibitors: Stattic and S3I-201 (Schust *et al*, 2006; Siddiquee *et al*, 2007). Both inhibitors significantly reduced the activation of STAT3 analyzed after incubation with apoptotic cancer cells for two hours. This treatment also impacted the stabilization of HIF-1α (Figure 2D–2G and Figure 2–Figure supplement 1). Similarly, these findings were observed in macrophages efferocytosing apoptotic Myc-CaP cells (Figure 2–Figure supplement 2). These findings strongly support the hypothesis that STAT3 activation is a critical signal that mediates the stabilization of HIF-1α by efferocytosis.

To further investigate these findings, a direct interaction between HIF-1α and p-STAT3 was evaluated in immunoprecipitation assays. Protein lysates collected from macrophages co-cultured with apoptotic prostate cancer RM1 cells were used to perform immunoprecipitation with HIF-1α-specific antibodies and the pull-down of p-STAT3 was evaluated by Western Blot. The blot showed a p-STAT3 band in the macrophage lysates from efferocytic samples which were immunoprecipitated with HIF-1α (Figure 2H). The p-STAT3 band was undetected in immunoprecipitated protein samples from non-efferocytic macrophages. These results strongly suggest the interaction between HIF-1α and p-STAT3 in efferocytic macrophages which support the hypothesis of STAT3 activation as a mechanism mediating the stabilization of HIF-1α.

### Efferocytosis stimulates the expression of pro-inflammatory MIF cytokine in macrophages

Accumulating experimental evidence suggests that efferocytosis of apoptotic cancer cells accelerates tumor progression and metastatic growth by fostering an inflammatory and immunosuppressive microenvironment (Elliott *et al*, 2017; Werfel & Cook, 2018). Single-cell data identified the negative regulation of the immune system process (GO: 0002683; related to the immunosuppressive response) as one of the GO terms upregulated in efferocytic macrophages. Using STRING, a database of known and predicted protein-protein interactions, we identified strong network association between this immune response and the biological process of hypoxia (GO: 0071456) (Figure 3A, GO gene list Table 2 and Table 3 in Supplementary data). Although not identified by single cell STAT3 was added because of its key role in the stabilization of HIF-1α as shown in Figure 2. Central nodes identified in this network are HIF-1α, Myc and STAT3 and the findings show direct or indirect interactions between hypoxia and the negative immune regulation processes. Furthermore, single cell analysis identified the cytokine macrophage migration inhibitory factor *Mif* as upregulated in efferocytic macrophages (p<10^-6^) (Figure 1F, Figure 3B and Figure 3–Figure supplement 1). *Mif* belongs to both GO: 0002683 and GO: 0071456 (Gene Ontology, 2021) and mediates both immunosuppression and inflammation and has been associated with increased tumorigenesis and disease progression in different cancer types including prostate cancer (Penticuff *et al*, 2019).

**Figure 3.**
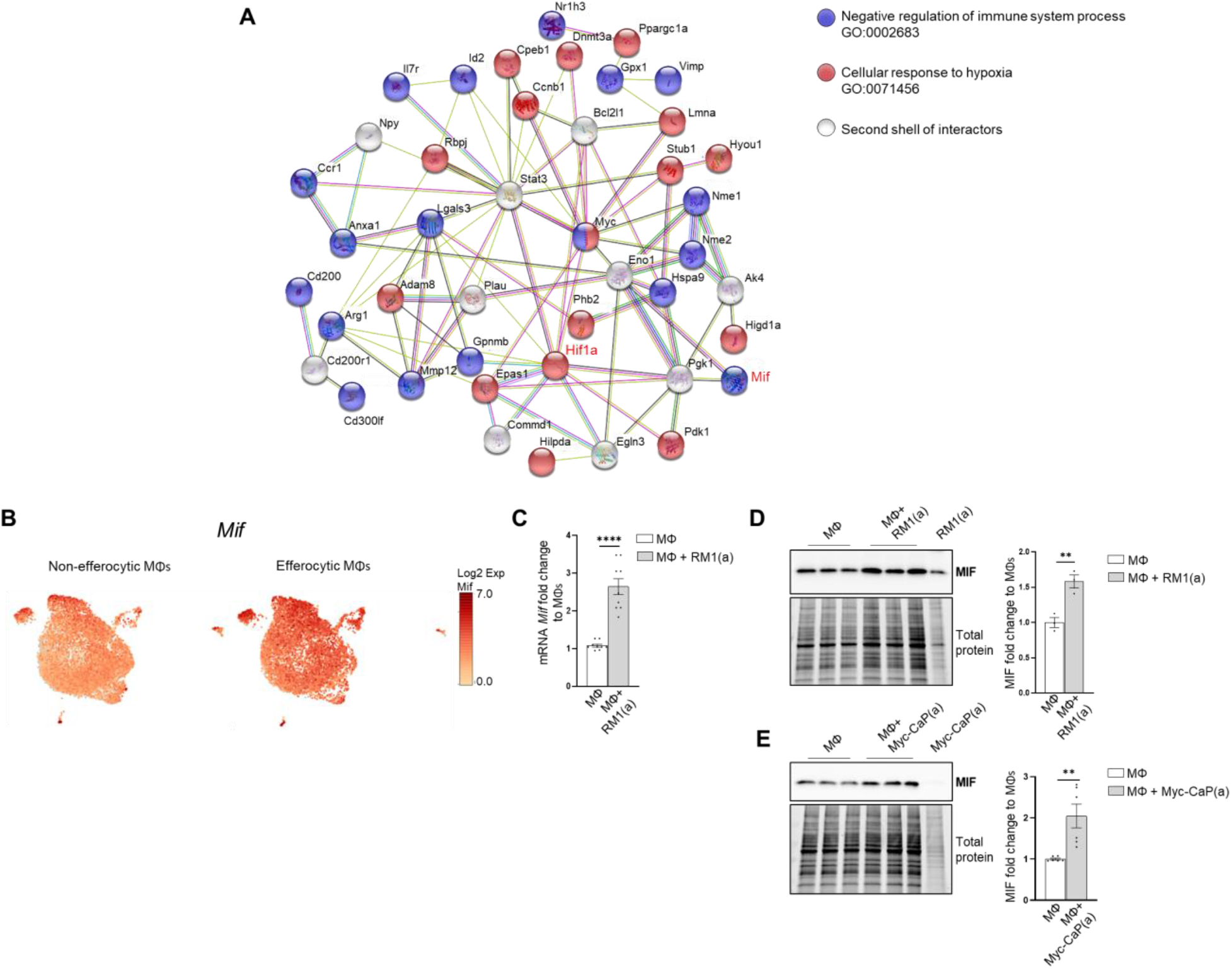
Macrophage efferocytosis induces MIF expression. **A.** Protein-protein interaction network between the immune pathway regulation and the hypoxia related pathway. Biological Process GO terms of selected genes upregulated by efferocytosis (GO:0002683, all genes; GO:0071456, 25 out of 33 genes). Bone marrow-derived macrophages were isolated from 4 week old C57BL/6J or FVB/NJ mice and co-cultured with apoptotic RM1(a) or Myc-CaP(a) cells for 16-18 hours. **B.** scRNA-Seq analysis plot shows *Mif* distribution in efferocytic and non-efferocytic clusters of macrophages. **C.** *Mif* mRNA expression in efferocytic macrophages assessed by RT-qPCR (n=9). Western blot analysis of protein lysates from **D.** C57BL/6J macrophages co-cultured with apoptotic RM1 prostate cancer cells (n=3) and **E.** FVB/NJ efferocytic and control macrophages using MIF antibody (n=6). Data plotted are mean ± SEM, **p < 0.01, ****p < 0.0001 (unpaired t-test). Additional results are shown in Figure 3–Figure supplement 1.

**Table 2.** Selected GO terms (Figure 1E) enriched in efferocytic bone marrow macrophages.

| GO term biological process | # of Genes in Term | # of GO term genes in upload list | Expected | Fold enrichment | Raw P-value | FDR value |
| --- | --- | --- | --- | --- | --- | --- |
| Regulation of innate immune response (GO:0045088) | 287 | 16 | 6.08 | 2.63 | 6.38E-04 | 3.89E-02 |
| Cellular response to hypoxia (GO:0071456) | 82 | 13 | 1.74 | 7.49 | 8.25E-08 | 2.90E-05 |
| Cellular response to decreased oxygen levels (GO:0036294) | 86 | 13 | 1.82 | 7.14 | 1.36E-07 | 4.20E-05 |
| Cellular response to oxygen levels (GO:0071453) | 101 | 14 | 2.14 | 6.55 | 1.18E-07 | 3.80E-05 |
| Translational initiation (GO:0006413) | 54 | 13 | 1.14 | 11.37 | 1.02E-09 | 9.44E-07 |
| Regulation of protein stability (GO:0031647) | 278 | 17 | 5.89 | 2.89 | 1.58E-04 | 1.44E-02 |
| Positive regulation of vasculature development (GO:1904018) | 200 | 17 | 4.23 | 4.02 | 3.13E-06 | 5.38E-04 |
| Regulation of response to wounding (GO:1903034) | 187 | 14 | 3.96 | 3.54 | 8.34E-05 | 8.73E-03 |
| Response to oxidative stress (GO:0006979) | 338 | 23 | 7.16 | 3.21 | 2.22E-06 | 4.27E-04 |
| Negative regulation of immune system process (GO:0002683) | 469 | 23 | 9.93 | 2.32 | 3.04E-04 | 2.35E-02 |

**Table 3.**
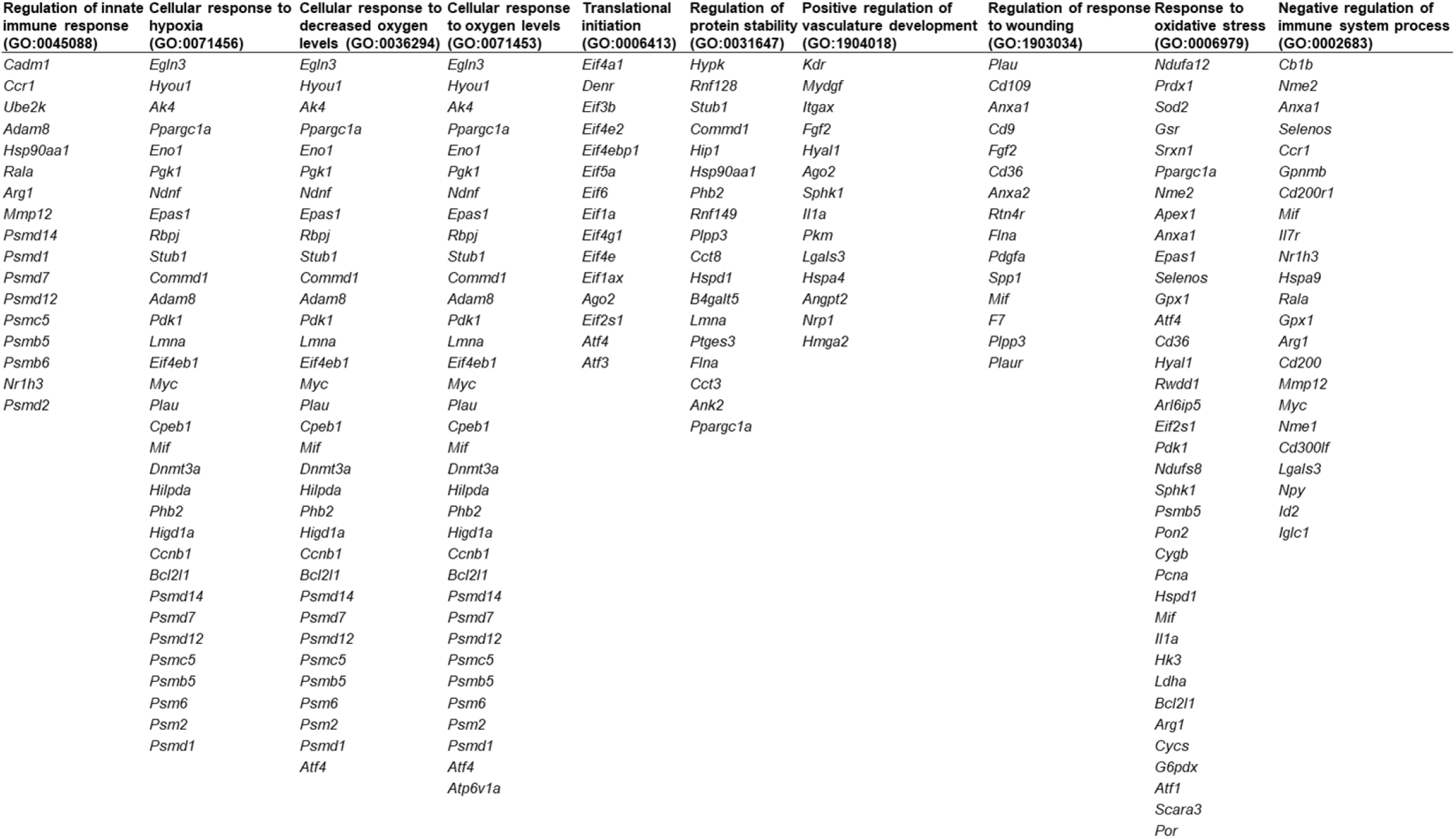
List of upregulated genes from GO terms shown in Figure 1E and Table 2.

MIF changes were investigated in the overall macrophage population co-cultured with apoptotic RM1 prostate cancer cells by RT-qPCR and Western blot. In correlation with single cell results both MIF mRNA (Figure 3C) and protein increased in bone macrophages efferocytosing apoptotic RM1 cells (Figure 3D). Similarly bone marrow macrophages isolated from FVB mice upregulated MIF protein upon efferocytosis of Myc-CaP prostate cancer cells (Figure 3E).

These findings indicate MIF is part of the signaling response of macrophages to the engulfing of apoptotic prostate cancer cells and suggests a network connection with the activation of hypoxia-related molecules by efferocytosis.

### HIF-1α mediates the expression of MIF cytokine in bone efferocytic macrophages

The implication of HIF-1α in tumor promoting inflammation, immunosuppression and metastasis has been documented in different cancer models in relation to hypoxia (Triner & Shah, 2016). We hypothesized that the stabilization of HIF-1α in bone macrophages by the clearance of apoptotic cancer cells induces the expression of key pro-inflammatory cytokines. This was investigated by crossing LysM-Cre mice with the HIF-1α ^flox/flox^ mice to obtain mutated HIF-1α myeloid lineage mutant mice (*Hif1a*^mut^). These mice create a null allele in the Cre-expressing cells (myeloid) lacking the exon 2 of *Hif1a* and were used to obtain *Hif1a*^mut^ bone macrophages. Relative *Hif1a* specific mRNAs were quantified with a primer/probe set corresponding to exon 2 of the mRNA. The qPCR analysis showed the upregulation of *Hif1a* mRNA in the efferocytic macrophages relative to control (Figure 4A). In support of the model a significant decrease in the *Hif1a* mRNA containing the exon 2 was observed in *Hif1a*^mut^ macrophages corresponding to control and efferocytic samples (Figure 4A). In addition, characterization of *Hif1a*^mut^ bone marrow derived macrophages by Western blot demonstrated a lower molecular weight in *Hif1a*^mut^ which corresponds with the deletion in the DNA binding domain encoded by the exon 2 (Figure 4C) by Cre-induced recombination which renders a non-functional *Hif1a*^mut^ protein. However even this mutant protein was stabilized by efferocytosis as shown by quantitative Western blot analyses, which suggests that STAT3-mediated stabilization is independent of HIF-1α binding to the chromatin (Figure 4C). Of note STAT3 activation remained unaffected in mutant macrophages (Figure 4–Figure supplement 1). Previous studies suggest a link between HIF-1α and MIF in different cell models, including macrophages (Alonso *et al*, 2019; Baugh *et al*, 2006). We investigated the expression of MIF and other pro-inflammatory factors as potential targets of HIF-1α. These included critical inflammatory cytokines previously found upregulated in efferocytic macrophages: CXCL1, CXCL5 and IL6 and CXCL4 (also known as platelet factor 4, *Pf4*) (Vaught *et al*, 2015). From the selected cytokines it was found that *Mif* and *Cxcl4* expression was significantly reduced in *Hif1a*^mut^ efferocytic macrophages relative to wild type (WT) after normalization to their specific control macrophages (Figure 4B).

**Figure 4.**
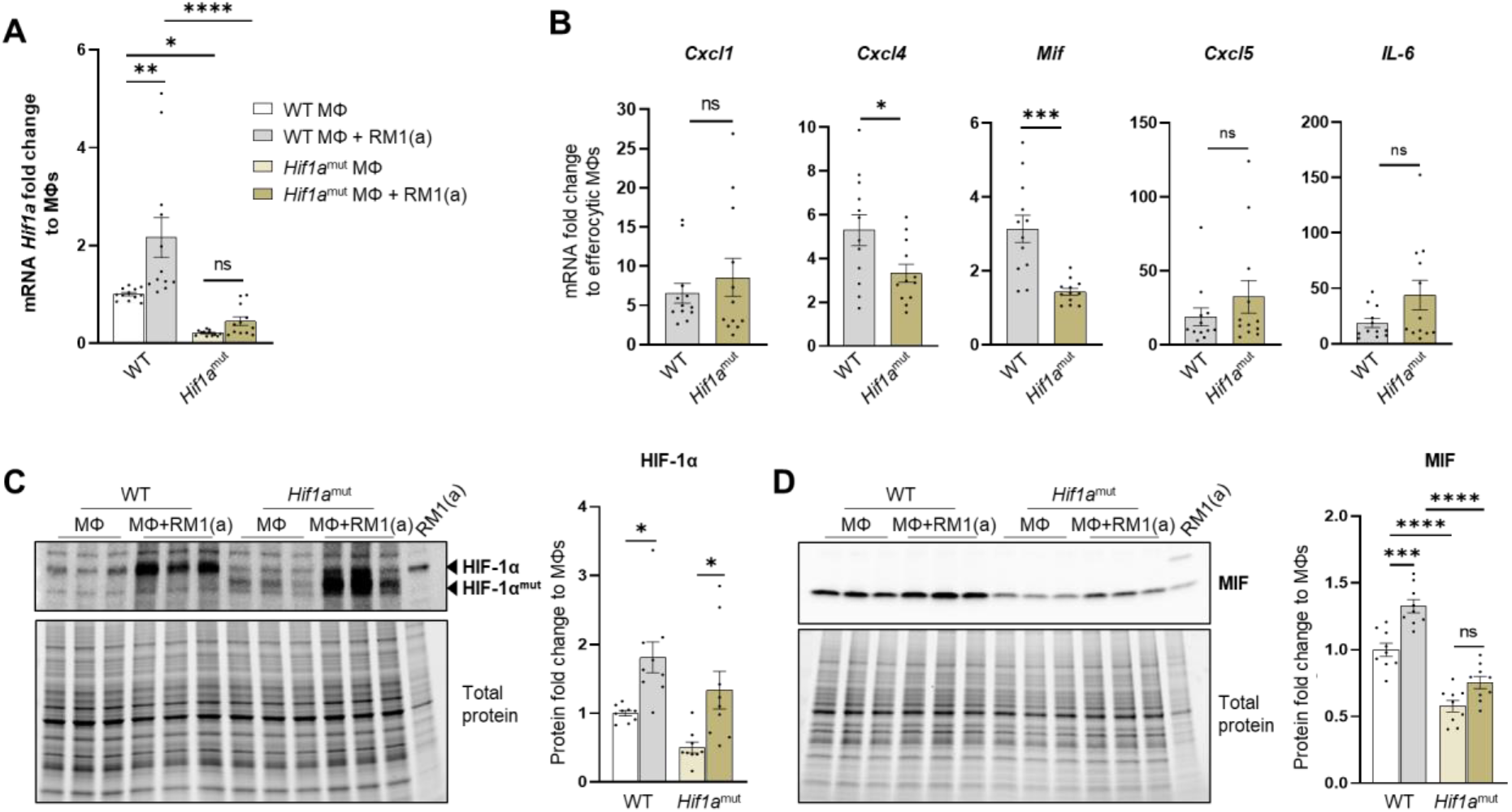
HIF-1α depletion in efferocytic macrophages reduces MIF expression. **A.** *Hif1a* mRNA expression levels in *Hif1a*^flox/flox^ and *Hif1a*^mut^ macrophages. **B.** mRNA from efferocytic and control macrophages from *Hif1a*^flox/flox^ and *Hif1a*^mut^ mice were analyzed by RT-qPCR for the specified inflammatory cytokine genes (n=12). **C. & D.** Protein lysates from efferocytic and control macrophages from *Hif1a*^flox/flox^ (WT) and *Hif1a*^mut^ mice were analyzed by Western blot with HIF-1α and MIF antibodies (n=9). Data plotted are mean ± SEM. *p < 0.05, ***p < 0.001, ****p < 0.0001, ns=not significant (Ordinary one-way ANOVA; Tukey’s multiple-comparisons test and unpaired t-test). Additional results are shown in Figure 4–Figure supplement 1 and 2.

Further analysis of MIF protein indicated a significant reduction in MIF protein expression and no upregulation was observed by efferocytosis in the *Hif1a*^mut^ macrophages relative to WT (Figure 4D). In contrast upregulation of MIF by efferocytosis was found in WT macrophages as previously shown in (Figure 3D–3E).

To address the specificity of HIF-1α in the control of MIF regulation, similar experiments were performed with *Epas1*-myeloid lineage-mutant (*Epas1*^mut^) mice. These mice were obtained by crossing LysM-Cre mice with the Epas1^flox/flox^ mice. The mutant mice expressed significantly lower levels of *Epas1* mRNA by RT-qPCR using the primer/probe set corresponding to the deleted exon2 (Figure 4–Figure supplement 2A). In addition, qPCR analysis of pro-inflammatory cytokines indicated a significant decrease in *Cxcl1* in the mutant mice, while no changes in *Mif* and *Cxcl4* (Figure 4–Figure supplement 2B), suggesting Epas1-specific regulation of pro-inflammatory cytokines, which is different than the observed for HIF-1α-mediated regulation in macrophages (Figure 4B). Quantitative protein analysis of mutant macrophages indicated no change by efferocytosis in HIF-1α stabilization in the *Epas1*^mut^ macrophages relative to WT control (Figure 4–Figure supplement 2C) and no change by efferocytosis in MIF expression when compared mutant versus WT (Figure 4–Figure supplement 2D) different from the results for efferocytic *Hif1a*^mut^ macrophages (Figure 4D). Furthermore, when analyzed relative to the control macrophages, the efferocytic *Epas1*^mut^ macrophages showed a significant increase in MIF (Figure 4– Figure supplement 2D), while no change was observed in efferocytic *Hif1a*^mut^ macrophages (Figure 4D). These results highlight the specificity in the regulation of MIF expression by HIF-1α relative to EPAS1, a similar hypoxia-inducible transcription factor.

HIF-1α-mediated MIF regulation was further investigated by using a HIF α-prolyl-hydroxylase inhibitor FG-4592 (also known as Roxadustat) (Hsieh *et al*, 2007). FG-4592 was used in efferocytosis assays, where a strong correlation was observed between the stabilization of HIF-1α and MIF protein expression. FG-4592 alone stabilized HIF-1α which correlated with up-regulation of MIF protein in non-efferocytic macrophages. Intriguingly, when the inhibitor was used in efferocytic macrophages a further increase in HIF-1α and MIF protein was observed (Figure 5A and 5B). RT-qPCR analysis showed a significant increase in *Mif* mRNA induced by FG-4592 relative to control however no further increase was observed in efferocytic macrophages (Figure 5C).

**Figure 5.**
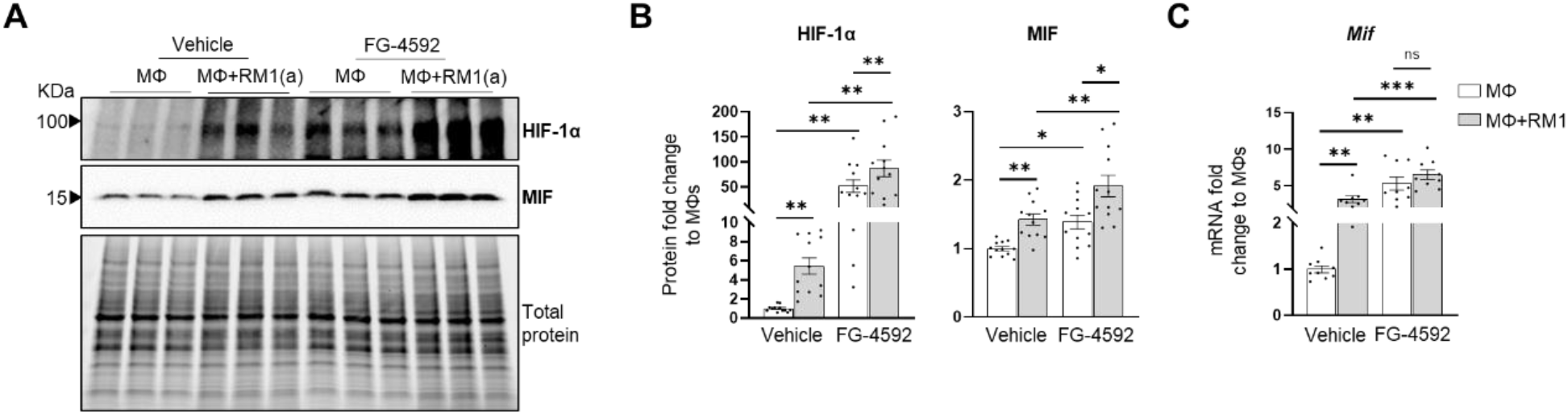
Efferocytic macrophages stabilize HIF-1α and induce MIF expression. Bone marrow-derived macrophages were isolated from C57BL/6J mice and co-cultured alone or with apoptotic RM1(a) cells and treated with FG-4592, HIF prolyl-hydroxylase inhibitor (10μM) or vehicle control for 16-18 hours. **A. & B.** Protein lysates from co-cultures were analyzed by Western blot with HIF-1α and MIF antibodies (n=12). **C.** mRNAs were isolated from co-cultures and analyzed by RT-qPCR for *Mif* expression (n=9). Graphs show the fold change relative to macrophage control for each group. Data plotted are mean ± SEM; *P<0.05, **P < 0.01, ***p < 0.001, ns=not significant (Repeated measures one-way ANOVA; Tukey’s multiple-comparisons test).

Altogether, these findings suggest that HIF-1α mediates the expression of MIF, where HIF-1α stabilization by efferocytosis or prolyl-hydroxylase inhibitor significantly upregulates MIF expression in bone macrophages.

### MIF activates inflammation in bone macrophages

CD74 is a critical receptor for MIF signal transduction in cells; however, CD74 lacks kinase activity and requires a complex formation with other co-receptors including CD44 and CXCR4 (Bernhagen *et al*, 2007; Leng *et al*., 2003; Shi *et al*., 2006). Results from single-cell data analyses identified a significant downregulation of CD74 in efferocytic macrophages as compared with non-efferocytic (Figure 6A, Figure 6–Figure supplement 1), while no significant differences were detected in the co-receptors CD44 or CXCR4. The downregulation of CD74 was also evident in the overall efferocytic macrophages relative to control by RT-qPCR analyses (Figure 6B).

**Figure 6.**
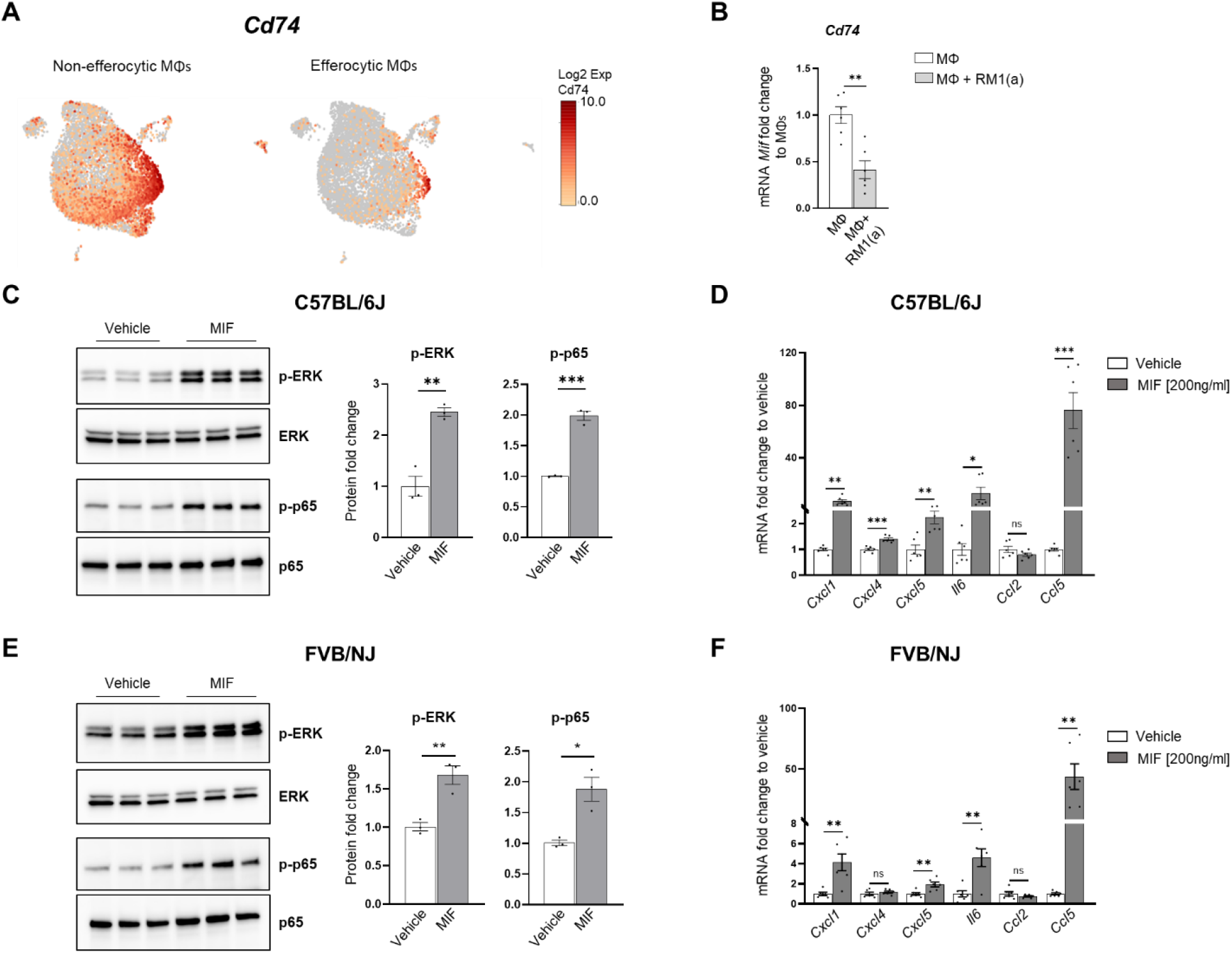
MIF induces a pro-inflammatory response in macrophages. Bone marrow-derived macrophages were isolated from C57BL/6J mice and treated with 200ng/ml of MIF protein or vehicle control during 2 hours for Western blot analysis and 8 hours for mRNA analysis. **A.** scRNA-Seq analysis plot shows *Cd74* distribution in efferocytic and non-efferocytic clusters of macrophages. **B.** *Cd74* mRNA expression in efferocytic macrophages assessed by RT-qPCR (n=6). **C.** Protein lysates from macrophages treated with MIF and vehicle control were analyzed by Western blot with total ERK, p-ERK, total p65 and p-p65 antibodies (n=3). **D.** mRNAs were isolated from co-cultures and analyzed by qPCR for the specified genes (n=6). Data are mean ± SEM; *p < 0.05, **p < 0.01, ***p < 0.001, ns=not significant (unpaired t-test). Additional results are shown in Figure 6–Figure supplement 1 and 2.

Although these results do not rule out a potential endocrine signaling, they suggest that MIF could induce a potent paracrine signaling in non-efferocytic macrophages and other cells. To evaluate the MIF-induced signaling in macrophages a purified recombinant MIF protein expressed in mammalian cells was used. Macrophages were incubated for 2 hours with MIF and signaling activation was analyzed by Western blot with specific phospho-peptide antibodies. A hallmark MIF transducing signal resulting in the activation (sustained phosphorylation) of the extracellular signal related kinase ERK1/2 MAPK (Calandra & Roger, 2003; Penticuff *et al*., 2019) was found highly upregulated in macrophages treated with recombinant MIF protein relative to control (Figure 6C and Figure 6–Figure supplement 2A). Furthermore, an increase in the critical inflammatory NF-kB signaling (phospho-p65) was observed in macrophages treated with MIF (Figure 6C and Figure 6–Figure supplement 2B).

This potent inflammation-transduced signaling pathway was further correlated with the increased expression in MIF activated macrophages of several pro-inflammatory cytokines including: *Ccl5*, *Cxcl5*, *Il6*, *Cxcl1* and *Cxcl4* (Figure 6D and Figure 6–Figure supplement 2B). Similar results were observed in FVB macrophages where MIF activated the expression of these cytokines, except for *Cxcl4* (Figure 6E). The activation of these cytokines has also been detected in macrophages engulfing prostate cancer cells and function to accelerate tumor growth in bone as previously demonstrated (Roca & McCauley, 2018). Altogether, these results suggest that the STAT3-HIF-1α-MIF is a potent signaling axis induced by efferocytosis of apoptotic cancer cells which may act via paracrine signaling to perpetuate inflammation.

## Discussion

Chronic inflammation has a major impact on cancer progression and metastasis in various organs (Coussens & Werb, 2002; Mantovani *et al*, 2008; Trinchieri, 2011). One of the inflammatory mechanisms that promotes tumor growth is the secretion of cytokines and chemokines by cancer and immune cells in the tumor microenvironment (de Visser *et al*, 2006; Quail & Joyce, 2013). Tumor-associated macrophages play a critical role in accelerating tumor progression in different cancer types (Aras & Zaidi, 2017; Duan & Luo, 2021). In bone, apoptotic cancer cell efferocytosis by macrophages generates a unique inflammatory milieu rich in cytokines that promote and support tumor progression (Mendoza-Reinoso *et al*., 2020; Roca *et al*., 2018). However, the molecular mechanisms that induce this tumor-promoting inflammatory response in bone marrow macrophages during the efferocytosis of apoptotic prostate cancer cells are still unclear.

Analysis of single-cell transcriptomics of sorted macrophages engulfing apoptotic prostate cancer RM1 cells (efferocytic, F4/80^+^CFSE^+^) versus sorted macrophages alone (non-efferocytic, F4/80^+^) by flow cytometry (Figure 1A) revealed two distinctive clusters of cells, each of them with a unique gene expression profile. Gene ontology term enrichment analysis of the upregulated genes in efferocytic macrophages identified several biological pathways including those directly related to macrophage functions such as response to wounding, innate immune response and regulation of vasculature development (Okabe & Medzhitov, 2016). Although the efferocytosis experiments in vitro were conducted under normal oxygen levels, the analysis identified GO-terms related to cell response to hypoxia or decreased oxygen levels in efferocytic macrophages, these terms included genes that are known targets of HIF-1α, a master transcriptional regulator of cell response to hypoxia (Cui *et al*, 2017). Similarly, hypoxia-independent HIF-1α expression has been observed in prostate cancer tumors, which correlates with recurrence following surgery or therapy, increased chemoresistance and accelerated metastatic progression, suggesting that alternative mechanisms of post-translational stabilization could lead to its accumulation and transcriptional activity in non-hypoxic environments (Ranasinghe *et al*, 2015; Ranasinghe *et al*, 2014). Here we found that efferocytic bone marrow macrophages promoted HIF-1α stabilization and induced a strong and sustained phosphorylation of STAT3 under normoxic conditions. Moreover, co-inmunoprecipitation experiments performed in this study demonstrated p-STAT3/HIF-1α interaction in efferocytic macrophages, which correlates with previous studies (Gray *et al*., 2005; Jung *et al*., 2008; Jung *et al*, 2005).

Previous studies have associated the expression of HIF-1α and its target genes with immunosuppressive functions in the tumor microenvironment (Chiavarina *et al*, 2010; Palazon *et al*, 2017). Here we identified a strong protein network association between the genes related to cellular response to hypoxia and negative regulation of the immune response in bone marrow efferocytic macrophages, suggesting that HIF-1α signaling in efferocytic macrophages may exert immunosuppressive functions in the tumor microenvironment. It has been reported that HIF-1α promotes the secretion of cytokines and chemokines such as CXC motif chemokine ligand 5 (CXCL5), CXC motif chemokine ligand 12 (CXCL12), chemokine ligand 28 (CCL28), and macrophage migration inhibitory factor (MIF) (Blaisdell *et al*., 2015; Du *et al*., 2008; Facciabene *et al*., 2011; Zhu *et al*., 2014). One of the genes included in the network analysis was *Mif*. MIF is a pro-inflammatory cytokine expressed by monocytes, macrophages, blood dendritic cells, B cells, neutrophils, eosinophils, mast cells, and basophils; and its expression is involved in both innate and adaptive immune processes, as well as in response to hypoxia. Several studies have shown that MIF mediates inflammatory processes such as sepsis and cancer (Bernhagen *et al*, 1993; O’Reilly *et al*, 2016).We demonstrated that MIF mRNA and protein levels were increased in bone marrow macrophages upon efferocytosis of apoptotic cancer cells. MIF is highly expressed in prostate cancer patients and its expression has been associated with higher severity and poor outcome (Meyer-Siegler *et al*, 2005). Also, it has been reported that HIF-1α regulates MIF secretion in breast cancer cells to promote tumor proliferation, angiogenesis and metastasis (Larsen *et al*., 2008). Interestingly, we found that HIF-1α but not HIF-2α depletion in bone marrow macrophages reduced the expression of the pro-tumorigenic inflammatory cytokines *Mif* and *Cxcl4* after being exposed to apoptotic prostate cancer cells when compared to wildtype bone marrow macrophages. Furthermore, HIF stabilization by a prolyl-hydroxylase inhibitor further stabilized HIF-1α and induced MIF mRNA and protein expression in efferocytic bone marrow macrophages. These results suggest that HIF-1α specifically regulates MIF expression in efferocytic macrophages, which may induce a chronic inflammatory response in the bone tumor microenvironment.

MIF signals through its main receptor CD74 and its co-receptors CD44, CXCR2, or CXCR4 (Bernhagen *et al*., 2007; Leng *et al*., 2003; Shi *et al*., 2006). MIF/CD74 activity promotes immunosuppressive signaling in macrophages and dendritic cells and inhibition of this signaling reestablishes the antitumor immune response in metastatic melanoma (Figueiredo *et al*, 2018; Tanese *et al*, 2015). Moreover, MIF/CD74 signaling also activates the NF-κB signaling pathway in chronic lymphocytic leukemia (Binsky *et al*, 2007; Gil-Yarom *et al*, 2017). CD74/CD44 activation by MIF is followed by phosphorylation of the proto-oncogene tyrosine-protein kinase (SRC), extracellular signal-related kinase 1/2 (ERK1/2), phosphoinositide 3-kinase (PI3K), and protein kinase B (AKT) (Gore *et al*, 2008; Leng *et al*., 2003; Lue *et al*, 2007; Shi *et al*., 2006). These kinases promote the activation of transcription factors such as nuclear factor-kappa B (NF-κB, p65) which induces the secretion of pro-inflammatory cytokines such as IL-6, IL-8, CCL2 and CCL5 (Abdul-Aziz *et al*., 2017; Binsky *et al*, 2007; Gregory *et al*, 2006; Johnson *et al*, 2018; Xu *et al*, 2008). Here we found that *Cd74* expression is downregulated in efferocytic bone marrow macrophages, suggesting a paracrine signaling in non-efferocytic macrophages. Recombinant MIF protein treatment of non-efferocytic macrophages activated the ERK1/2 and the NF-κB (p65) pathways and increased the expression of pro-inflammatory cytokines such as CXCL1, CXCL5, IL-6 and CCL5.

Altogether, these findings reveal a new regulatory mechanism of HIF-1α in macrophages during efferocytosis of apoptotic cancer cells where the p-STAT3/HIF-1α/MIF signaling pathway induces chronic inflammation in the tumor bone microenvironment and promotes the expression of pro-inflammatory cytokines (Figure 7). Since enhanced cell death inevitably occurs during tumor growth and is increased with many cancer therapies our results suggest this pathway becomes highly activated in bone metastatic patients undergoing chemo or radiation therapies and may contribute to increased inflammation and accelerated cancer growth.

**Figure 7.**
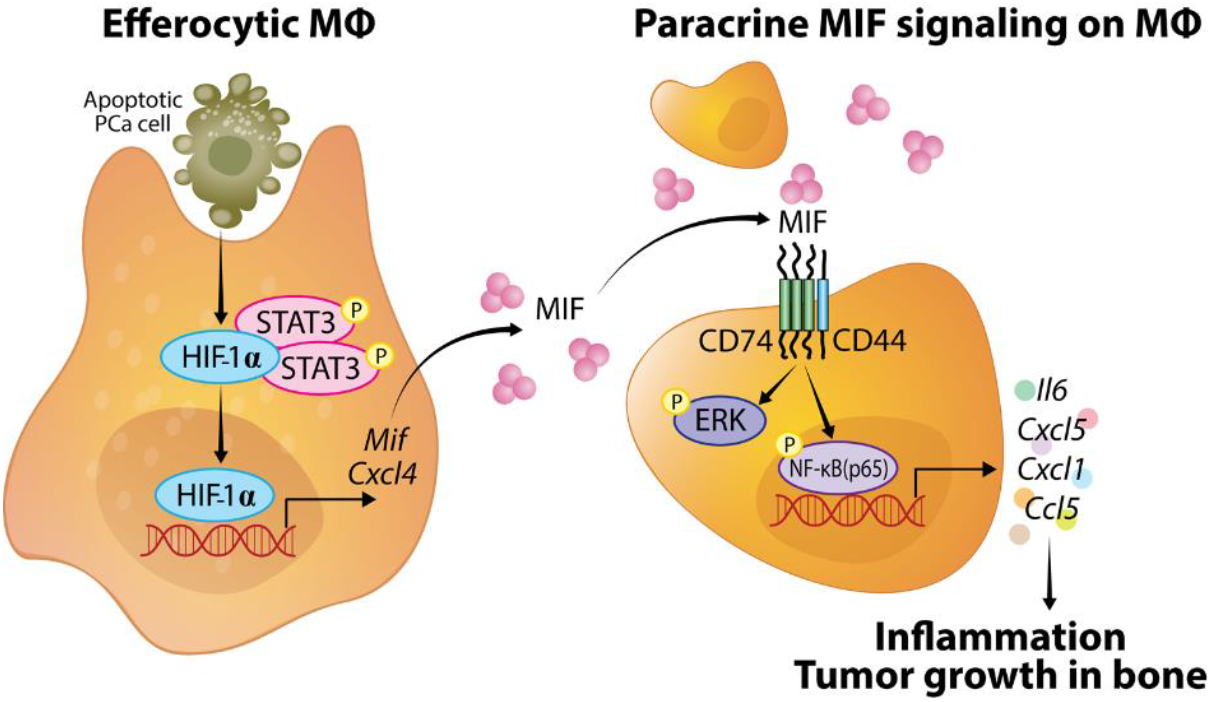
HIF-1α signaling during bone macrophage efferocytosis of apoptotic cancer cells. Bone macrophage engulfment of apoptotic prostate cancer cells promotes HIF-1α stability by its interaction with p-STAT3. Once HIF-1α is stabilized it is translocated to the nucleus to initiate the transcription and secretion of *Mif* and *Cxcl4*. Secreted MIF binds CD74/CD44 receptor complex of neighboring macrophages and activates ERK1/2 and p65 to induce the production of pro-inflammatory cytokines in the tumor microenvironment to support tumor growth in bone.

## Material and Methods

### Key resources table

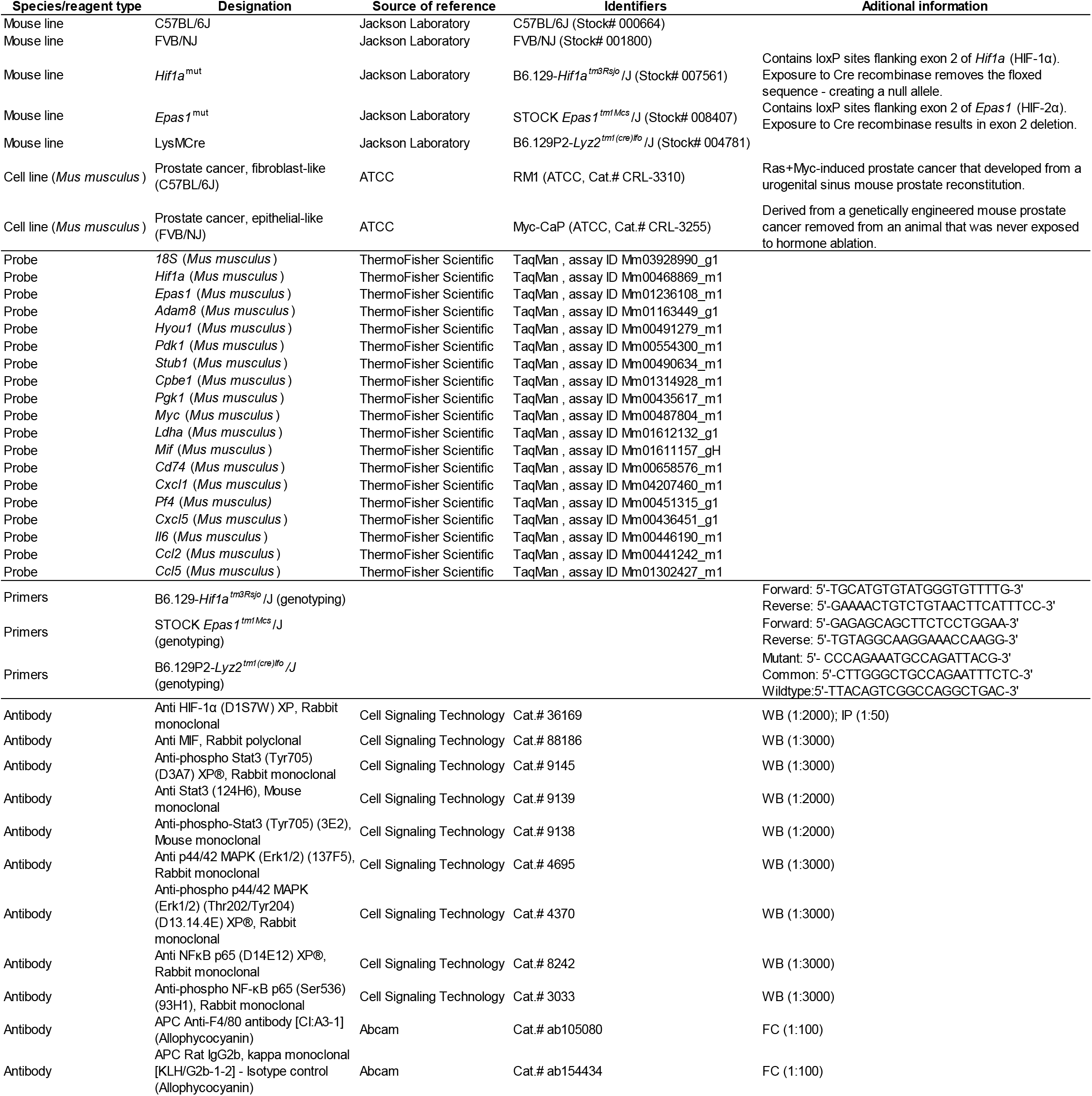

### Animals and cell lines

All animal experiments were performed with approval from the University of Michigan Institutional Animal Care and Use Committee. Immunocompetent C57BL/6J, FVB/NJ, B6.129P2-*Lyz2^tm1(cre)Ifo^*/J (LysMCre), B6.129-*Hif1a^tm3Rsjo^*/J (HIF-1α^flox^) (Ryan *et al*, 2000) and *Epas1^tm1Mcs^*/J (HIF-2α^flox^) (Gruber *et al*, 2007) mice were purchased from the Jackson Laboratory (Bar Harbor, ME, USA). The HIF-1α^flox^ and HIF-2α^flox^ mice were crossed consecutively with LysMCre mice to achieve the Hif1a^flox/flox^-LysMCre^+/-^ (*Hif1a*^mut^) and Epas1^flox/flox^-LysMCre^+/-^ (*Epas1*^mut^) mice that exhibit HIF-1α and HIF-2α inactivation in myeloid cells including macrophages. HIF-1α^flox/flox^ and HIF-2α^flox/flox^ mice were used as experimental controls.

RM1 is a (Ras+Myc)-induced prostate cancer cell line developed in C57BL/6J mice and was a gift from Timothy C. Thompson (Baylor College of Medicine, Houston, TX, USA) (Baley *et al*, 1995; Thompson *et al*, 1989). Myc-CaP prostate cancer cell line is derived from a prostate carcinoma from a Hi-Myc FVB/NJ mice and was donated by Russell Taichman and Frank Cackowski (University of Michigan, Ann Arbor, MI, USA) (Watson *et al*, 2005). Both cell lines were cultured on RPMI 1640 media containing 10% fetal bovine serum (FBS) and grown at 37°C with ambient O2 and 5% CO_2_.

### Murine efferocytosis in vitro model

Bone marrow-derived macrophages (MΦs) were isolated from 4–6 week old male C57BL/6J, FVB/NJ, *Hif1a*^mut^, *Epas1*^mut^, HIF-1α^flox^ and HIF-2α^flox^ mice by flushing the femur and tibia with minimum essential medium eagle - alpha modification (αMEM) supplemented with L-glutamine, antibiotic-antimycotic 1× and 10% fetal bovine serum (FBS). Macrophages were cultured in αMEM (L-glutamine, antibiotic-antimycotic 1×, 10% FBS) in the presence of macrophage colony stimulating factor (M-CSF) (30 ng/mL, #315-02, Peprotech, Rocky Hill, NJ, USA). After three days in culture, macrophages were plated independently at 2 × 10^6^ cells/well in αMEM (L-glutamine, antibiotic-antimycotic 1×, 0.25% FBS) for co-culture experiments. RM1 and Myc-CaP cells were exposed to UV light for 30 min to induce apoptosis. Apoptotic (a) cells (>90% trypan blue incorporation) were co-cultured with macrophages at a 1:1 ratio in αMEM (L-glutamine, 0.25% FBS) for 16–18 h. Macrophages from mutant mice were compared with those from respective littermate controls.

Prolyl hydroxylase inhibition in efferocytic and non-efferocytic macrophages was performed using 10μM Roxadustat (FG-4592) (Cayman Chemical, 15294) for 16–18 hours. STAT3 inhibition in efferocytic and non-efferocytic macrophages was performed using 5μM Stattic (Cayman Chemical, 14590) or 60μM S3I-201 (Millipore Sigma, SML0330) for 10 minutes, then media was replaced and incubated for an additional 2 hours. Macrophages alone were treated with 200ng/ml of recombinant MIF protein (SinoBiological, 50066-M08H) during 2 hours for Western blot analysis and 8 hours for RNA expression analysis.

### Single-cell library preparation and RNA sequencing

A modified murine efferocytosis *in vitro* model was used. Apoptotic RM1 cells were labeled with CellTrace™ CFSE (ThermoFisher Scientific, C34554), and then co-cultured with macrophages for 16–18 h. Efferocytic and non-efferocytic macrophages were collected and incubated in fluorescence-activated cell sorter (FACS) staining buffer (phosphate buffered saline-1X, 0.2% bovine serum albumin). F4/80 antibody and isotype control were added and incubated for 1 h at 4 °C. F4/80^+^ only (non-efferocytic macrophages) and F4/80^+^CFSE^+^ (efferocytic macrophages) were sorted using a BD FACSAriaTM III (BD biosciences, San Jose, CA, USA). Antibody information is available in Key resources table.

The single cell scRNA-seq libraries were prepared at the University of Michigan Advanced Genomics Core using the 10X Genomics Chromium Next GEM Single Cell 3’ Kit v3.1 (part number 1000268) following the manufacturer’s protocol. Cell suspensions were diluted to target a recovery of 10,000 cells per sample. The libraries were run on an Agilent TapeStation 4200 (part number G2991BA) for library quality control before sequencing. Libraries were sequenced at a depth of 50,000 reads/cell on a NovaSeq6000 with the following run configuration: Read 1 - 150 cycles; i7 index read - 8 cycles; Read 2 - 150 cycles.

### Single-cell RNA-sequencing analysis and visualization

The sequenced data was processed using the 10X Genomics CellRanger software suite v3.0.0. Briefly, fastq files from each of the samples were mapped to the Mouse genome mm10 and genes were counted using CellRanger software and the STAR aligner (Dobin *et al*, 2013). The barcode-gene matrices were further analyzed using the Seurat R package (v3.1) (Butler *et al*, 2018). Following standard practices to remove low-quality cells, cells that expressed less than 200 genes or less than 1,000 transcripts, or had greater than 10% mitochondrial genes were filtered from the datasets (Supplementary Table 1). For genes, only the top 5,000 variable genes were included for downstream analysis. Samples were then normalized and integrated according to the Seurat suggested pipeline. To reduce the dimensionality of the samples, we first performed a principal component analysis (PCA). The number of principle components for further downstream applications were 20, and UMAP was employed for final dimensionality reduction and visualization of the data.

### Differential Expression and Gene Ontology Analysis

Differential expression analysis was conducted using the DESingle R package (Miao *et al*, 2018). Genes with a false discovery rate adjusted p-value < 0.05 were considered differentially expressed. For pathway analysis, we used PANTHER analysis (Mi *et al*., 2013; Thomas *et al*., 2003) with the gene ontology database (Ashburner *et al*., 2000; Gene Ontology, 2021). Only genes differentially expressed and up-regulated in the efferocytic macrophages in both experiments were included in the GSEA analysis.

### Western blot analysis and co-immunoprecipitation assay

Whole cell lysate was extracted in Cell Lysis Buffer 1X (Cell signaling, 9803) containing 1X protease and phosphatase inhibitor cocktail (ThermoFisher Scientific, 78440). Estimation of protein concentration was done using Bradford assay (BioRad, 5000006). Samples were diluted using 1X Laemmli Sample Buffer (4X stock, BioRad, 1610747) with 10% β-Mercaptoethanol (Millipore Sigma, M3148). Protein lysates were separated using 4-20% Mini-PROTEAN^®^ TGX Stain-Free^™^ gels (BioRad, 4568096) and transferred to PVDF membrane using the Trans-Blot Turbo RTA kit (BioRad, 1704272). The membrane was blocked with 5% milk in 1X TBS-0.1% Tween for 1 hour at room temperature, then incubated with primary antibodies in 5% BSA during overnight at 4°C. Secondary antibody was diluted in 5% milk in 1X TBS-0.1% Tween. For co-immunoprecipitation assays macrophage and apoptotic prostate cancer cell co-cultures were lysed on ice with 1% Triton X-100 in 1X PBS with 1X protease and phosphatase inhibitor cocktail. Whole cell lysates were immunoprecipitated using HIF-1α antibody and protein A-magnetic beads (Cell Signaling Technology, 73778) during overnight incubation at 4°C. Binding and washing were performed in the same lysis buffer followed by immunoblotting with appropriate antibodies. Blots were developed using SuperSignal^™^ West Femto Maximum Sensitivity Substrate (ThermoFisher Scientific, 34095). Protein gels used for protein normalization and blots were imaged using the ChemiDoc™ MP Imaging System (BioRad, 12003154). Antibody information is available in Methods - Key resources table.

### RT-qPCR

Cells were harvested using RNeasy Mini Kit (Qiagen, 74106) RNA was eluted with nuclease-free water. The RNA was quantified using a NanoDrop 2000 (Thermo Scientific) and cDNA was synthetized 1μg of RNA per 20 μl reaction mixture using High-Capacity cDNA Reverse Transcription Kit (ThermoFisher Scientific, 4368814). RT-qPCR was performed using TaqMan^®^ probes and Gene Expression qPCR Assays TaqMan Gene Express (ThermoFisher Scientific, 4369016) with 40 cycles on an ABI PRISM 7700 (Applied Biosystems, Foster City, CA, USA). The analysis was performed using 2-ΔΔCT method (Schmittgen & Livak, 2008). TaqMan^®^ probes information is available in Key resources table.

### Statistics

Statistical analyses were performed using GraphPad Prism 9 (GraphPad Software, version 9.1.0, San Diego, CA, USA) using ordinary and repeated measures one-way analysis of variance (ANOVA) with Tukey’s multiple-comparisons, and unpaired t-tests analyses with significance of p < 0.05.

## Data availability

Raw sequencing data for both experiments: Experiment 1 (GSM5466517/roca4, non-efferocytic and GSM5466518/roca5, efferocytic). Experiment 2 (GSM5466519/roca6, non-efferocytic and GSM5466520/roca7, efferocytic) are deposited in the NCBI Sequence Read Archive (SRA) and can be accessed from the NCBI Gene Expression Omnibus (GEO, Series Accession: GSE180638).

## Acknowledgements

The authors would like to thank the University of Michigan Advanced Genomics Core for the single-cell processing and sequencing of the samples, and the Rogel Cancer Center Single Cell Analysis Core for further analysis of the single-cell data included is this study.

## Supplementary data

**Figure 2–Figure supplement 1.**
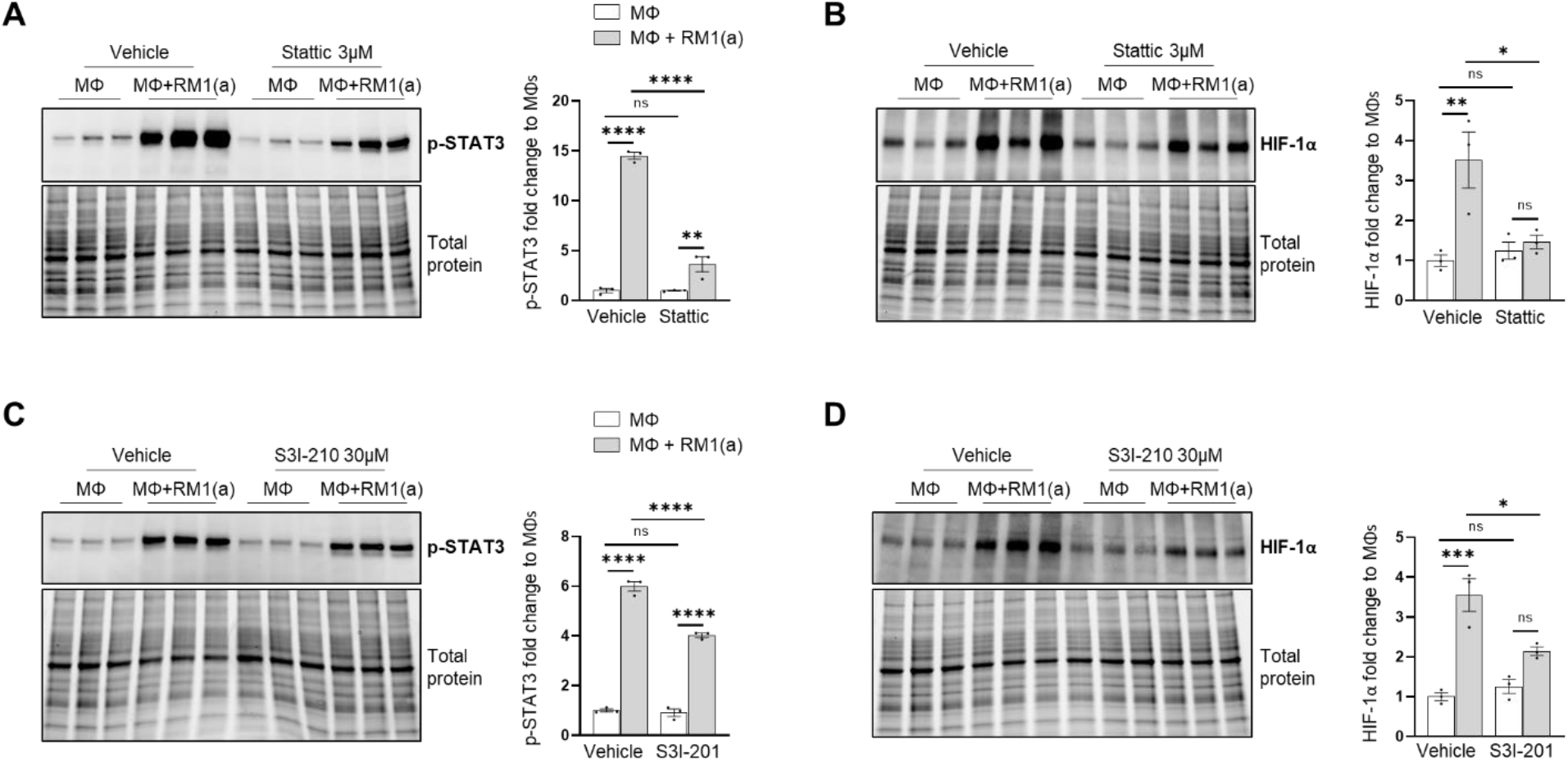
STAT3 inhibition (Stattic and S3I-201) in C57BL/6J in bone marrow macrophages efferocytosing apoptotic prostate cancer RM1 cells (n=3 per group).

**Figure 2–Figure supplement 2.**
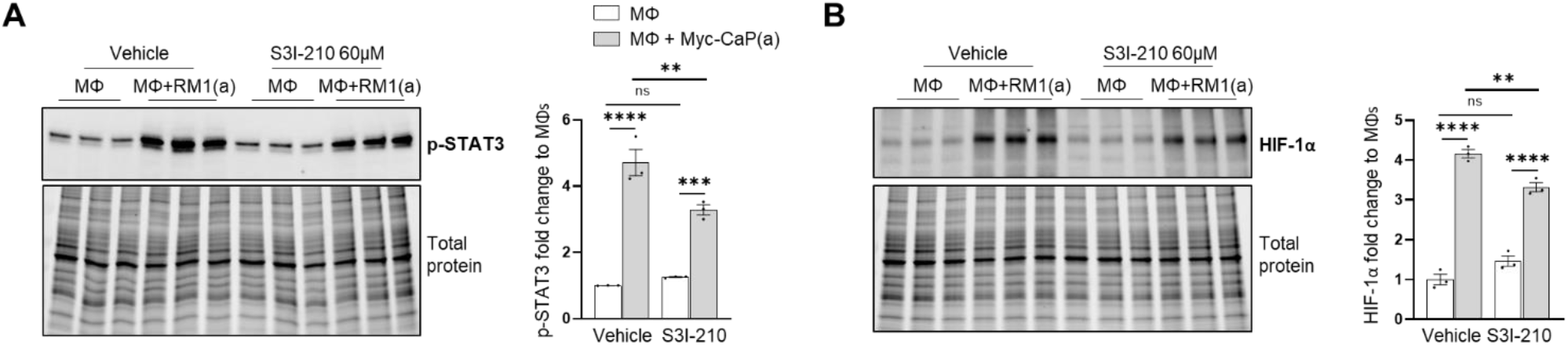
STAT3 inhibition (S3I-201) in FVB/NJ bone marrow macrophages efferocytosing apoptotic prostate cancer Myc-CaP cells (n=3 per group).

**Figure 3–Figure supplement 1.**
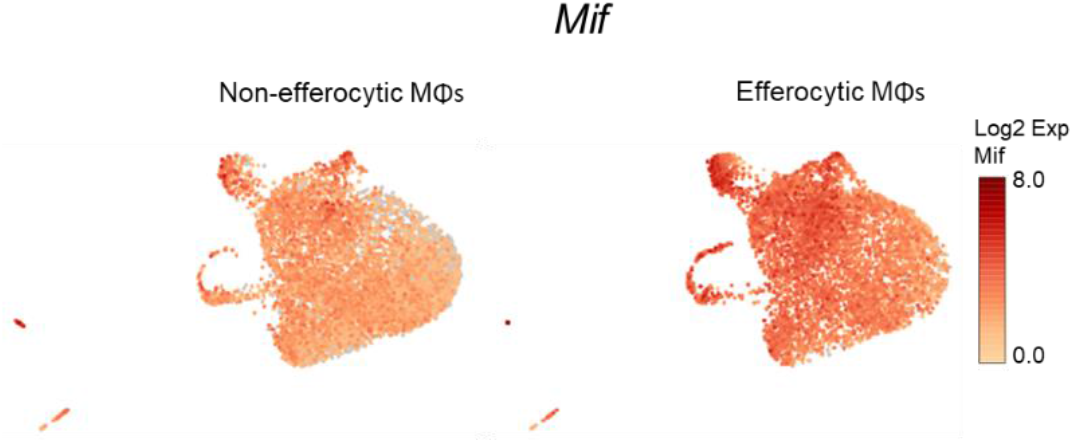
Single-cell plots of *Mif* expression in efferocytic macrophages (corresponding to Exp. 1)

**Figure 4–Figure supplement 1.**
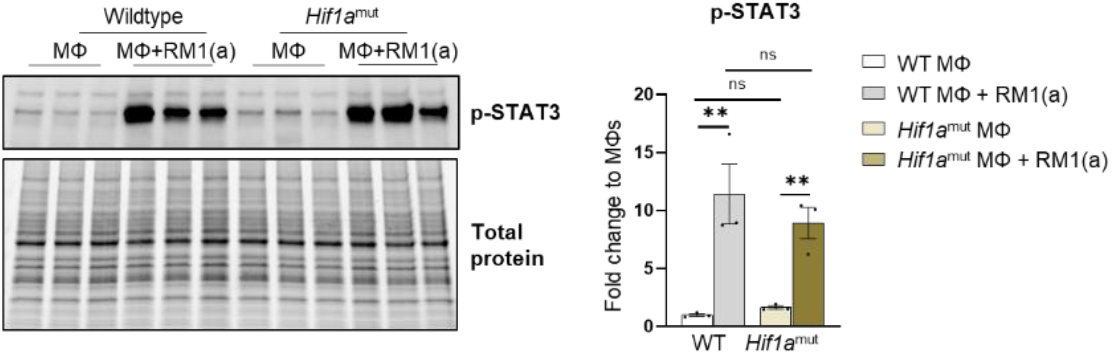
p-STAT3 expression in efferocytic *Hif1a*^mut^ macrophages compared to control macrophages (n=3).

**Figure 4–Figure supplement 2.**
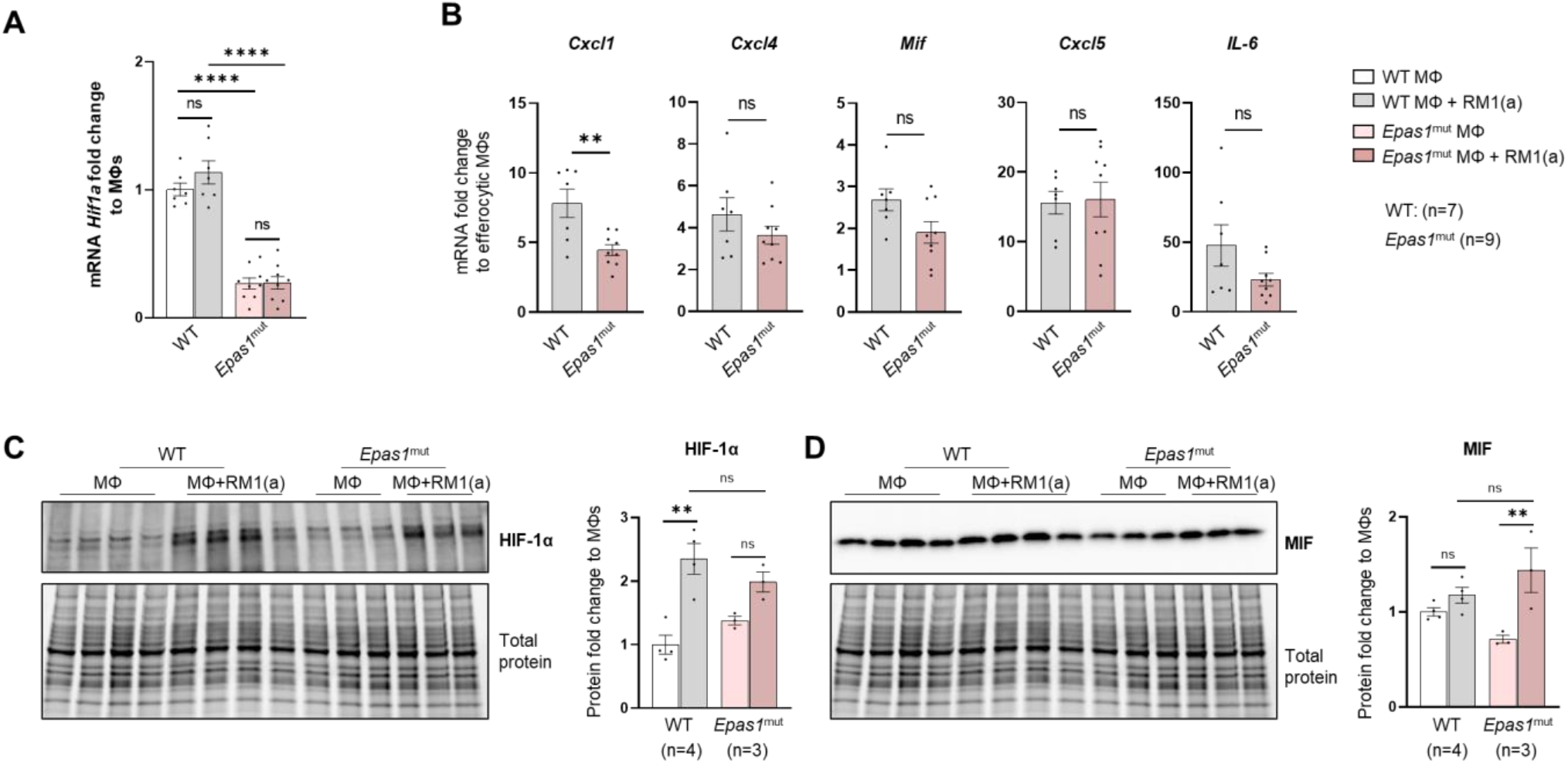
HIF-2α depletion (*Epas1*^mut^) effect in efferocytic macrophages.

**Figure 6–Figure supplement 1.**
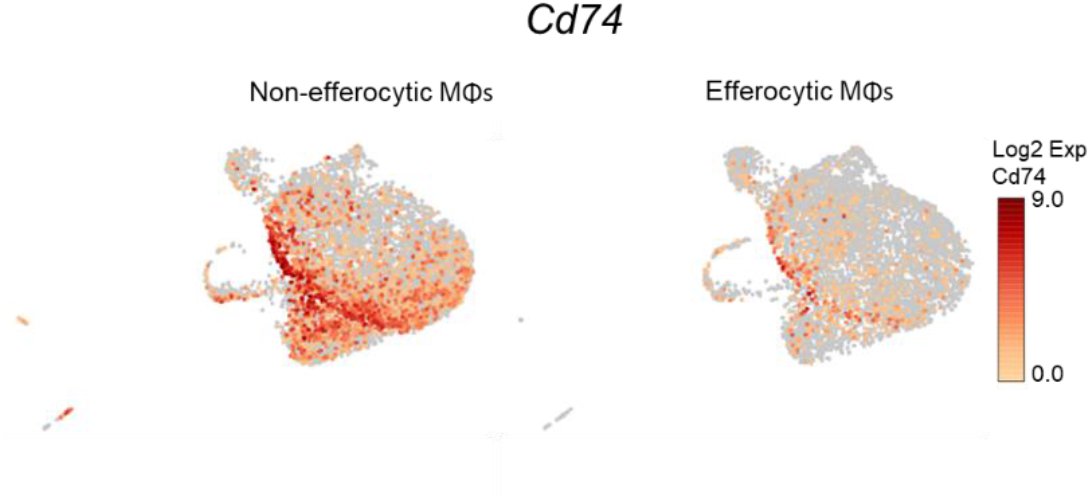
Single-cell plots of *Cd74* expression in efferocytic macrophages (corresponding to Exp. 1).

**Figure 6–Figure supplement 2.**
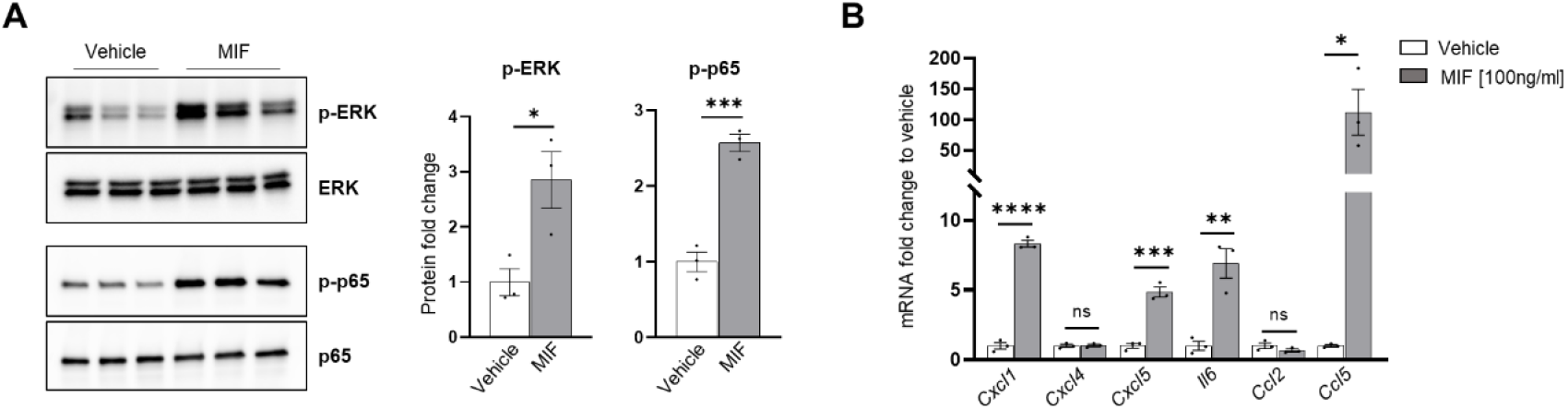
C57BL/6J bone marrow macrophages treatment with MIF (100ng/ml, n=3 per group).

